# Expectation triggers a change-like EEG response without acoustic change

**DOI:** 10.64898/2026.04.13.718093

**Authors:** Haoxuan Xu, Wanshun Wen, Yinru Yu, Ishrat Mehmood, Zhouyou Dai, Maida Mariam, Baorong Zhang, Xiongjie Yu

**Affiliations:** Department of Anesthesia, Women’s Hospital, Zhejiang University School of Medicine, Hangzhou, China; Department of Anesthesiology, Shanghai Tenth People’s Hospital, Tongji University School of Medicine, Shanghai, China; Zhejiang Provincial Key Laboratory of Precision Diagnosis and Therapy for Major Gynecological Diseases, Women’s Hospital, Zhejiang University School of Medicine, Hangzhou, China; College of Biomedical Engineering and Instrument Science, Zhejiang University, Hangzhou, China; Key Laboratory for Biomedical Engineering of Ministry of Education, Zhejiang University, Hangzhou, China; Department of Neurology of the Second Affiliated Hospital of Zhejiang University School of Medicine, Hangzhou, China; Hangzhou Entel Foreign Language School, Hangzhou, Zhejiang, China

**Keywords:** Predictive coding, Temporal uncertainty, Electroencephalography (EEG), Change detection, Click trains

## Abstract

Predictive processing theories posit that the auditory system continuously generates expectations and updates internal models when incoming evidence violates those expectations. Whether such updating can be expressed as a temporally precise, change-like neural response at a context-defined moment during continuous stimulation––without any physical change––has remained unclear. Using a transitional click-train paradigm in human EEG, we show that listeners reliably detect subtle midpoint transitions when click timing is fixed, whereas temporal uncertainty markedly weakens this sensory-driven detection. Strikingly, even in a physically unchanged stimulus, variable click timing elicits a robust response time-locked to the nominal midpoint. This endogenous change-like signal is stimulus-statistics dependent––absent in the fixed-timing no-change control––temporally confined to the early post-change window, and strongly state dependent, being amplified during active change detection relative to passive listening. Moreover, within identical no-change trials, the response tracks subjective change reports. Together, these findings show that temporal uncertainty and task engagement jointly gate predictive updating in continuous auditory streams, providing a compact assay to dissociate sensory-driven change responses from context- and state-dependent predictive signals.

## Introduction

Listening is fundamentally predictive. In natural scenes, the auditory system rarely waits for sounds to “arrive” before interpreting them; instead, it continuously anticipates structure unfolding in time—when a footstep should land, when a syllable should end, when a pattern should turn. Predictive-coding and related inference frameworks formalize this intuition: the brain maintains generative models of sensory causes, evaluates incoming evidence against those models, and updates them when predictions fail ^1–3^. One of the most robust population-level signatures of violated expectation is mismatch negativity (MMN), elicited when a deviant breaks an established regularity ^4–6^. At the neuronal level, stimulus-specific adaptation (SSA) can likewise be interpreted as a predictive response: repeated inputs become expected and are down-weighted, whereas rare or deviant events elicit relatively preserved responses, consistent with local prediction-error signaling ^7–14^. Together, these observations support the modern view that MMN-like signals reflect inference over sequence structure and the consequences of violated expectations, rather than a simple reaction to novelty ^4–6,15^.

A striking implication of this framework is that neural updating can be aligned to expected moments even when there is little—or no—corresponding acoustic energy. In omission paradigms, the absence of an anticipated sound evokes time-locked cortical activity, demonstrating that prediction-related signals can be anchored to internal timing rather than to stimulus onset or physical change ^16,17^. Hierarchical designs further show that prediction errors are sensitive to higher-order structure: when the brain infers that a deviant should occur at a particular position, omitting that expected deviant can elicit stronger responses than omitting a standard, consistent with multi-level predictions and errors ^15,18^. Related work using nonlinguistic structured stimuli has likewise revealed hierarchical temporal processing across the primate thalamocortical system ^19^. Moreover, expectations need not be driven solely by external stimulus statistics. Evidence from “phantom” or internally generated percepts suggests that endogenous perceptual states can establish predictions that yield deviance-like responses when violated ^20^.

Despite these advances, two gaps limit our understanding of predictive updating during continuous stimulation. First, omission-related responses are most often studied in event-based or strongly rhythmic contexts, where entrainment and phase-reset can contribute to time-locked activity, complicating the question of whether prediction-related update signals can be expressed without discrete events and without overt rhythmic scaffolding ^16,17^. Second, expectation-induced effects (including “phantom deviant”-like phenomena) are rarely tested when the critical update time is embedded within continuous input and defined primarily by task context—for example, an anticipated midpoint transition—rather than by the occurrence (or omission) of discrete tokens ^15,20^. Thus, it remains unclear whether change-related neural machinery can be recruited as a temporally precise, context-defined update signal even when the stimulus contains no physical change, and how such updating depends on stimulus temporal statistics and behavioral engagement.

To address these questions, we used a transitional click-train paradigm in which two continuous segments are concatenated and a nominal change point is fixed at the midpoint. Within the same dataset ^21^, we identified a midpoint-locked predictive change-like response that emerges without any physical change: in the no-change condition with variable click timing around the same mean (Irreg_4–4_), EEG showed a significant response aligned to the nominal transition, which was context dependent, enhanced by active detection, and absent in the no-change control with fixed click timing (Reg_4-4_). We propose that increased temporal uncertainty reduces the precision of bottom-up evidence and increases reliance on top-down priors about *when* change is likely, enabling an internally timed update signal even when the stimulus remains constant.

## Results

We established an experimental context in which participants could form a strong temporal expectation about when a change might occur. Each trial consisted of a 2-s click train composed of two consecutive 1-s segments (Train 1 and Train 2), with a fixed nominal change point at 1 s (Fig. 1A).

**Figure 1.**
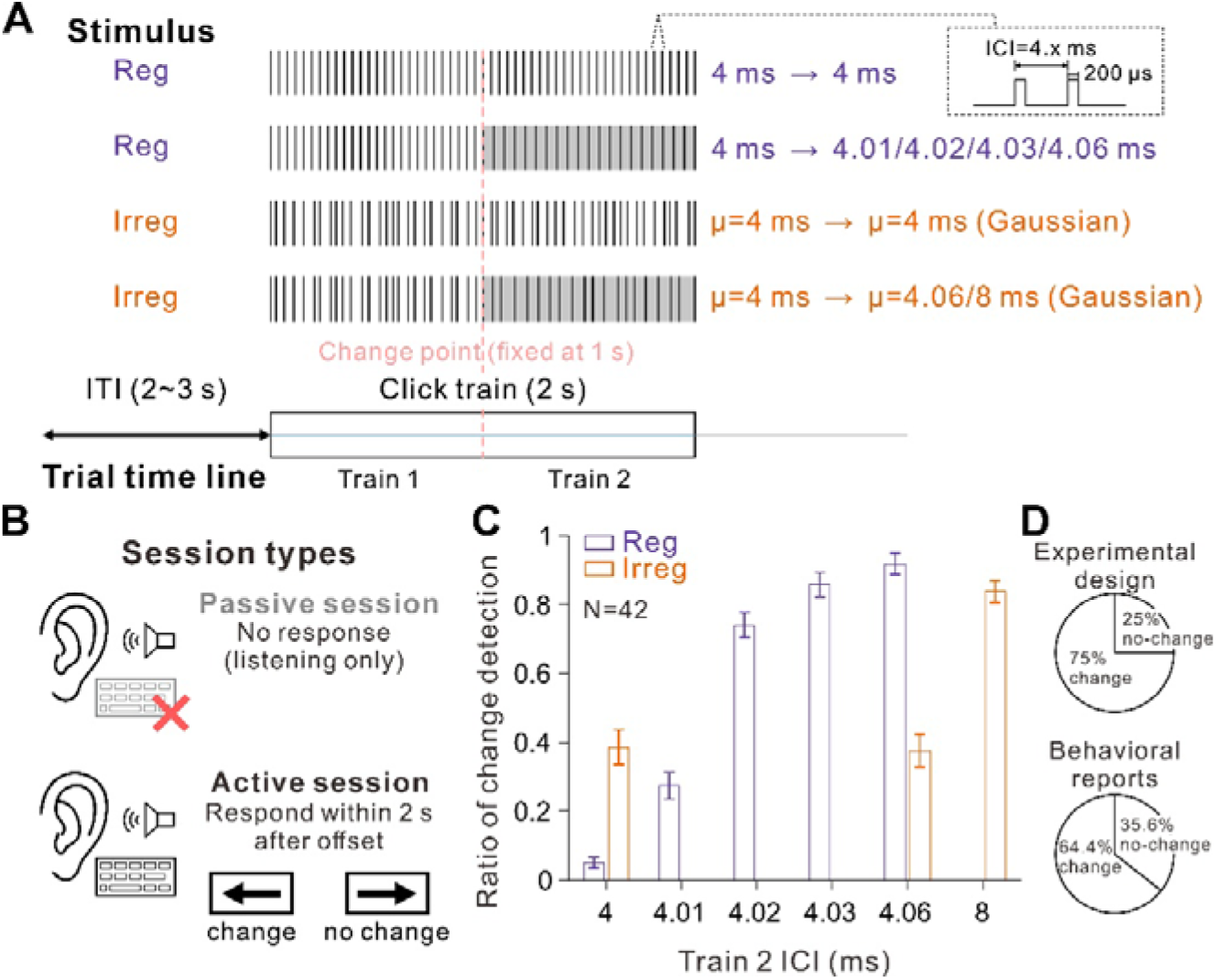
Experimental design and behavioral results. **(A) Stimuli and trial structure.** Each trial consisted of a 2-s click train comprising two consecutive 1-s segments (Train 1 and Train 2), with a fixed nominal change point at 1 s. Eight stimulus types were presented with equal probability (12.5% each): Reg_4–4_ (no change), Reg_4–4.01/4.02/4.03/4.06_ (changes in Train 2 click timing), Irreg_4–4_ (no change), and Irreg_4–4.06_ and Irreg_4–8_ (changes in Train 2 mean timing). In the Irreg conditions, click timing was drawn from a Gaussian distribution with fixed mean values. Click duration was 200 μs and the inter-trial interval (ITI) ranged from 2 to 3 s. **(B) Sessions.** Participants first completed a passive listening session without behavioral report, followed by an active change-detection session in which they reported “change” or “no change” via key press within 2 s after stimulus offset. **(C) Behavior.** Change-detection ratio in the active session plotted as a function of Train 2 timing. Purple bars denote Reg conditions and orange bars denote Irreg conditions. Data are mean ± s.e.m. across participants (N = 42). **(D) Stimulus proportions and behavioral reports.** Top: programmed proportions of no-change versus change trials. Bottom: observed proportions of “no change” versus “change” reports pooled across trials.

Eight stimulus types were randomly interleaved with equal probability (12.5% each): regular no-change (Reg_4–4_), regular change (Reg_4–4.01_, Reg_4–4.02_, Reg_4–4.03_, Reg_4–4.06_), irregular no-change (Irreg_4–4_), and irregular change (Irreg_4–4.06_ and Irreg_4–8_) (Fig. 1A). In the irregular conditions, click timing was sampled from Gaussian distributions with fixed mean values, and all trials used identical timing of the nominal change point (Fig. 1A). Because six of the eight trial types contained a change and all changes—when present—occurred at the fixed midpoint, participants were predisposed to monitor the 1-s timepoint for a potential transition.

Participants (N = 42) completed two sessions in sequence: a passive listening session without any behavioral report, followed by an active change-detection session in which they indicated “change” or “no change” within 2 s after stimulus offset (Fig. 1B). In the active session, change reports depended strongly on Train 2 timing and on stimulus statistics (Fig. 1C). For fixed-timing trains, the probability of reporting a change increased monotonically with the magnitude of the Train 2 timing shift (Reg_4–4.01/4.02/4.03/4.06_ relative to Reg_4–4_), indicating high sensitivity to small timing differences when timing was stable (Fig. 1C). By contrast, when click timing was variable, change reports were elevated even in the no-change condition (Irreg_4–4_) and did not reliably separate Irreg_4–4.06_ from Irreg_4–4_ (*p* = 0.68, Wilcoxon signed rank test), whereas a large timing shift (Irreg_4–8_) was readily detected (Fig. 1C). Together, these results confirm that temporal variability increases subjective “change-likeness” and reduces sensitivity to small timing differences, while preserving detection for large transitions.

The trial composition and behavioral judgments further quantify this change-expectant context. By design, 75% of trials contained a physical change (six change types out of eight), and 25% were no-change trials (Fig. 1D, upper). Behaviorally, participants reported “change” on 64.4% of trials and “no change” on 35.6% of trials (Fig. 1D, lower), indicating a global bias toward change reports that is consistent with the change-prevalent stimulus context and sets the stage for expectation-driven midpoint-locked responses in EEG.

We next tested whether a change-like EEG response could emerge without any physical transition, time-locked to the nominal change point at 1 s. In the active session, we compared the two no-change stimuli, Reg_4–4_ and Irreg_4–4_ (Fig. 2A–C).

**Figure 2.**
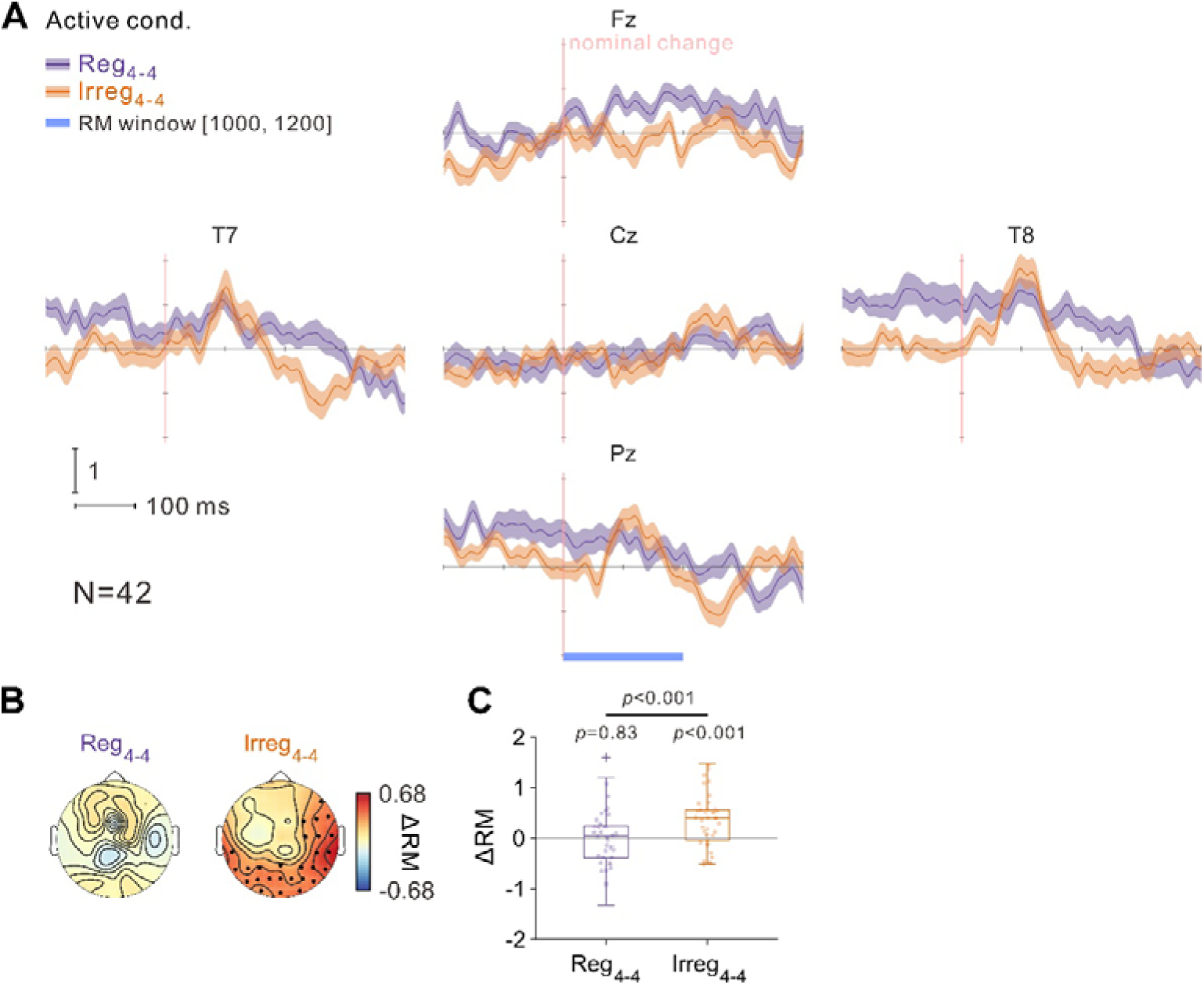
Predictive change-like responses in the no-change stimulus with variable click timing. **(A)** Grand-average ERPs (N = 42) from five example channels (Fz, T7, Cz, T8, and Pz) for the no-change stimuli Reg_4–4_ (purple) and Irreg_4–4_ (orange) in the active session; shaded bands indicate ± s.e.m. The vertical pink line marks the nominal change point (1 s). The blue bar indicates the RM window (1000–1200 ms post-onset). **(B)** Scalp distributions of ΔRM for Reg_4–4_ and Irreg_4–4_. The color scale indicates ΔRM magnitude (warm colors, positive ΔRM; cool colors, negative ΔRM). Black dots denote electrodes with ΔRM significantly greater than zero (one-sample *t*-test against zero, FDR-corrected *p* < 0.05). **(C)** Participant-level mean ΔRM averaged across the predefined temporal-parietal-occipital ROI. *p* values above each condition indicate one-sample *t*-tests against zero; the horizontal comparison indicates the paired *t*-test between conditions.

Across five example channels, Irreg_4–4_ showed a pronounced change-locked modulation—most evident at T7, T8, and Pz—whereas Reg_4–4_ exhibited little or no response at the same timepoint (Fig. 2A). We quantified this effect using ΔRM computed within the predefined RM window (1000–1200 ms post-onset; blue bar, Fig. 2A). Here, RM denotes the RMS amplitude of the ERP within the RM window (baseline-corrected using a pre-change interval), and ΔRM denotes the corresponding baseline-normalized response magnitude used for between-condition comparisons (see Methods). Scalp topographies revealed significant ΔRM for Irreg_4–4_ (black dots), primarily over temporal–parietal–occipital regions, whereas Reg_4–4_ showed no significant change-like effects (Fig. 2B). At the participant level, mean ΔRM averaged across the predefined temporal–parietal–occipital ROI (23 channels) was significantly larger for Irreg_4–4_ than for Reg_4–4_ (Fig. 2C), demonstrating a predictive change-like response expressed in the absence of a physical change and selectively enhanced when click timing is variable.

To confirm that the paradigm also yields canonical sensory-driven change responses when a physical change is present, we compared transition and no-change conditions within each stimulus class. In the active session, Reg_4–4.06_ elicited a robust change response relative to Reg_4–4_, with clear between-condition differences in example channels (Supplementary Fig. 1A), widespread significant ΔRM across electrodes (Supplementary Fig. 1B), and a strong participant-level separation in average ΔRM (Supplementary Fig. 1C). In contrast, the corresponding contrast under variable timing (Irreg_4–4.06_ vs Irreg_4–4_) was substantially weaker, though still detectable at the group level in the active session (Supplementary Fig. 1D–F). These results establish a positive control: physical transitions produce strong change responses when timing is stable, but the sensory-driven change response is reduced when timing is variable.

We then tested whether the predictive change-like response depends on task engagement by comparing Irreg_4–4_ between passive and active sessions (Fig. 3A–C).

**Figure 3.**
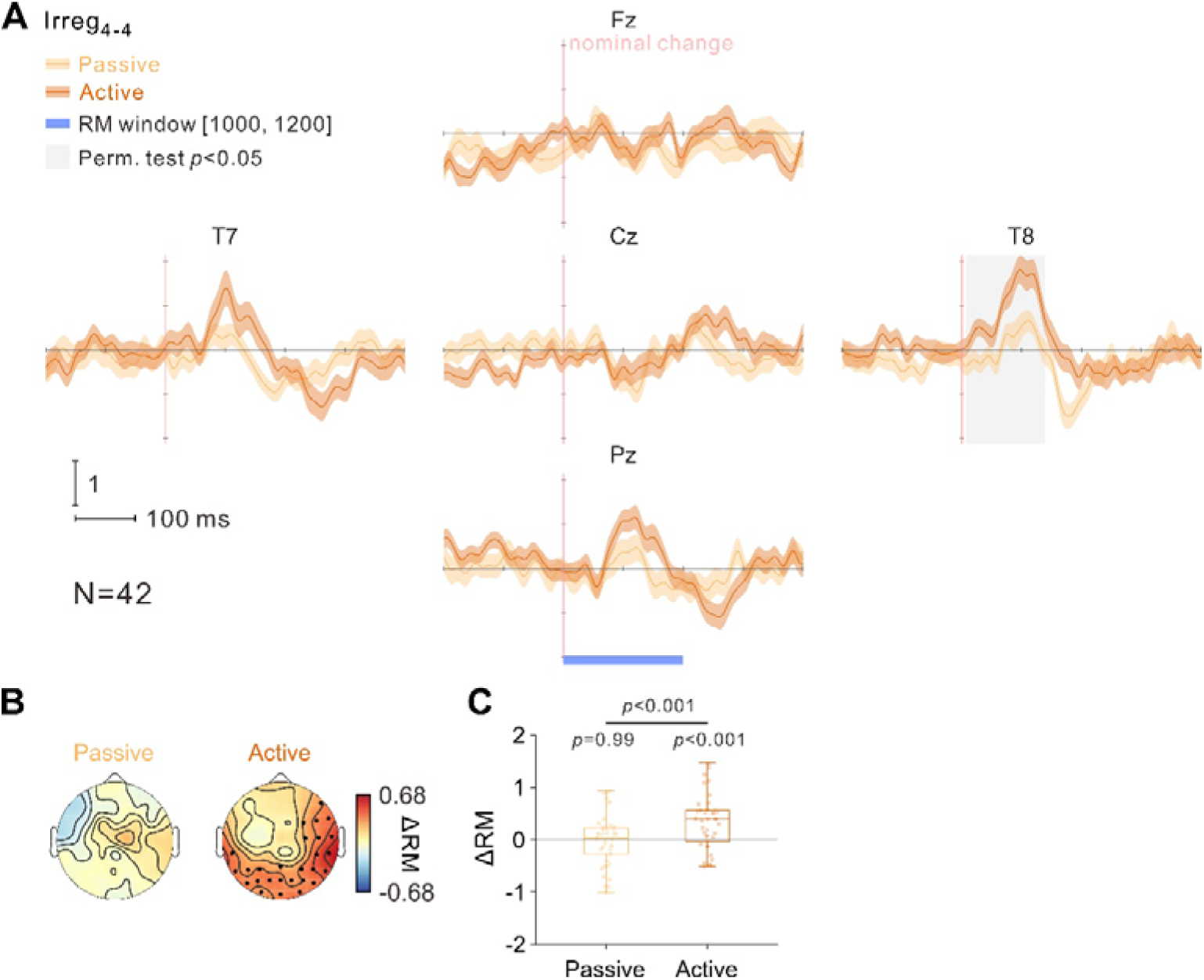
Task-dependent enhancement of the predictive change-like response in Irreg_4–4_. **(A)** Grand-average ERPs (N = 42) from five example channels (Fz, T7, Cz, T8, and Pz) for Irreg_4–4_ in the passive (light orange) and active (orange) sessions; shaded bands indicate ± s.e.m. The vertical pink line marks the nominal change point (1 s). The blue bar indicates the RM window (1000–1200 ms post-onset). Vertical gray bar indicates significant difference between conditions (*p* < 0.05, permutation test). **(B)** Scalp distributions of ΔRM for Irreg_4–4_ in the passive and active sessions. The color scale indicates ΔRM magnitude (warm colors, positive ΔRM; cool colors, negative ΔRM). Black dots denote electrodes with ΔRM significantly greater than zero (one-sample *t*-test against zero, FDR-corrected *p* < 0.05). **(C)** Participant-level mean ΔRM averaged across the predefined temporal-parietal-occipital ROI. *p* values above each condition indicate one-sample *t*-tests against zero; the horizontal comparison indicates the paired *t*-test between sessions.

In the same example channels, the change-locked modulation for Irreg_4–4_ was markedly stronger during active detection than during passive listening (Fig. 3A), and permutation testing identified a significant time window in channel T8 (gray shading, Fig. 3A). This task dependence was also evident in ΔRM topographies (Fig. 3B) and in participant-level average ΔRM across channels, which was significantly higher in the active than in the passive session (Fig. 3C). Consistent with this, in the passive session the stable-timing transition remained robust (*p* < 0.001, paired t-test, Reg_4–4.06_ > Reg_4–4_; Supplementary Fig. 2A–C), whereas the variable-timing transition contrast was not significant (Irreg_4–4.06_ vs Irreg_4–4_; Supplementary Fig. 2D–F). Thus, active engagement selectively enhanced the predictive change-like response in the variable-timing no-change condition while revealing a broader dissociation between stable-timing and variable-timing sensory-driven change responses.

Two additional controls strengthened the interpretation that the Irreg_4–4_ effect reflects a change-like response rather than a generic midpoint artifact. First, the fixed-timing no-change control (Reg_4–4_) showed no predictive change-like response and no significant active–passive difference in ΔRM (Supplementary Fig. 3A–C), arguing against a general session effect or nonspecific midpoint segmentation. Second, a time-window comparison aligned to the nominal change point demonstrated temporal specificity of the Irreg_4–4_ response (Fig. 4).

**Figure 4.**
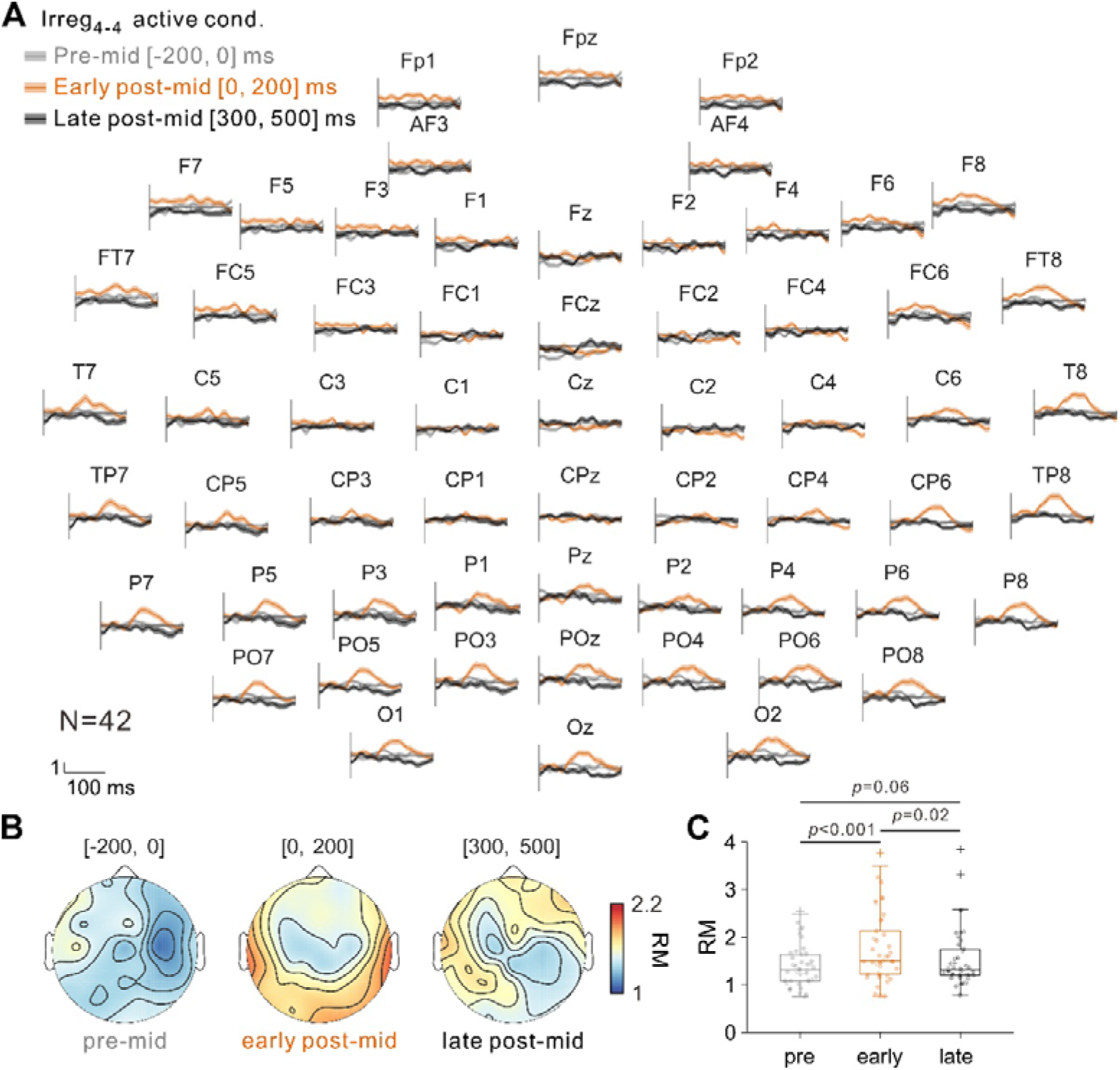
Temporal specificity of the predictive change-like response. **(A)** ERPs for Irreg_4–4_ in the active session (N = 42) are shown for all scalp electrodes. For each electrode, three waveform segments aligned to the nominal middle change point (1 s) are overlaid: pre-mid (gray, −200 to 0 ms), early post-mid (orange, 0 to 200 ms), and late post-mid (black, 300 to 500 ms). **(B)** Scalp distributions of RM for the same three time windows. The color scale indicates RM magnitude. **(C)** Participant-level mean RM averaged across the predefined temporal-parietal-occipital ROI for each time window. Horizontal comparisons indicate paired *t*-tests between time windows (pre vs early post: *p* < 0.001; early post vs late post: *p* = 0.02; pre vs late post: *p* = 0.06).

When waveform segments were overlaid across electrodes, a clear deflection was evident in the early post-mid interval (0–200 ms), whereas the pre-mid (−200–0 ms) and late post-mid (300–500 ms) intervals showed no comparable response (Fig. 4A). This pattern was mirrored in RM topographies (Fig. 4B) and in ROI-averaged RM, which was significantly larger in the early post-mid window than in the pre-mid and late post-mid windows (Fig. 4C). Together, these controls indicate that the predictive change-like response is specifically time-locked to the expected change point, rather than reflecting a broad fluctuation across the change-centered epoch.

Finally, we tested whether the predictive change-like response in Irreg_4–4_ covaries with participants’ subjective reports on physically identical no-change trials. We therefore separated Irreg_4–4_ trials in the active session according to behavioral report into trials judged as “change” and trials judged as “no change” (Fig. 5A).

**Figure 5.**
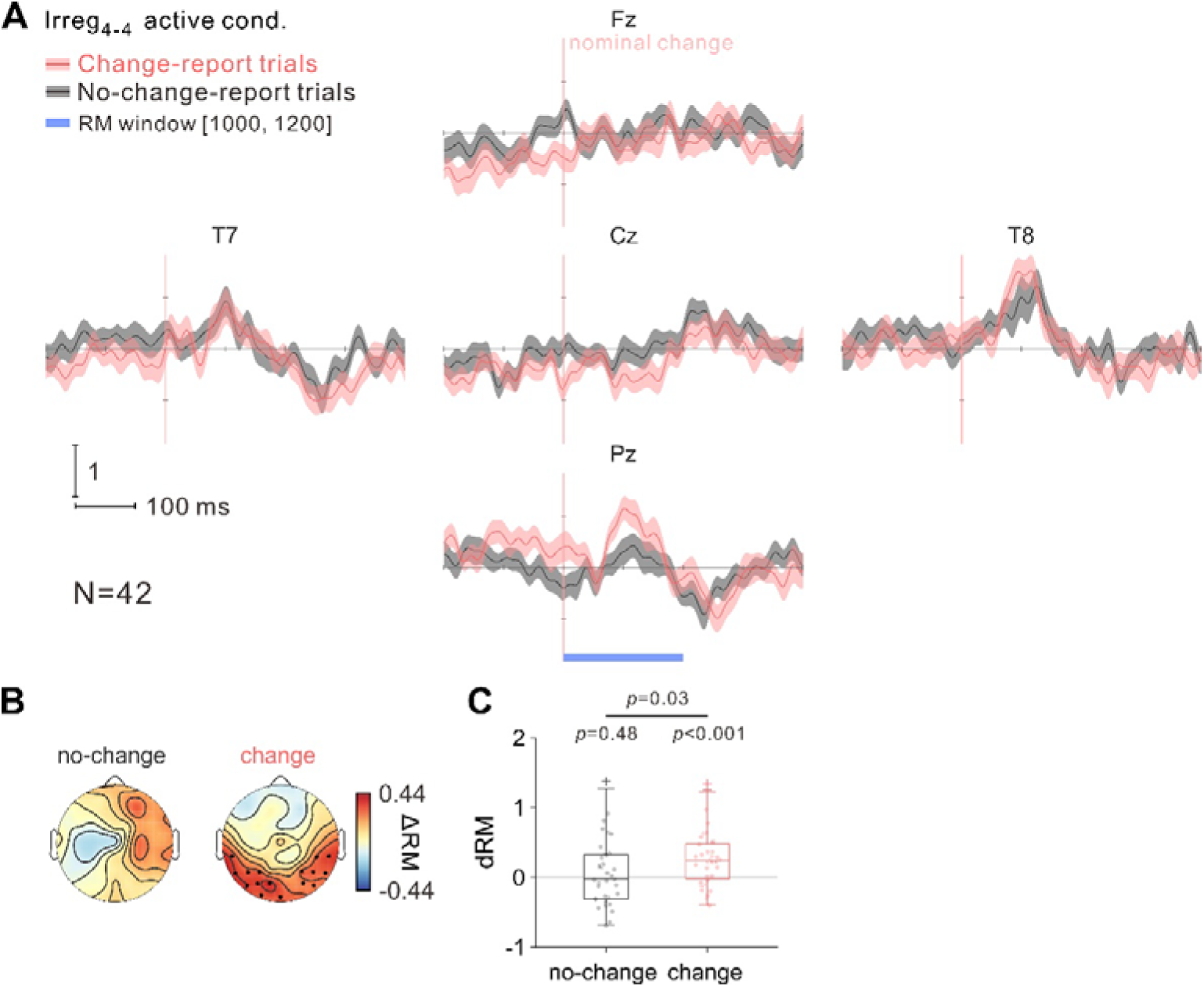
Predictive change-like responses track subjective change reports. **(A)** Grand-average ERPs (N = 42) from five example channels (Fz, T7, Cz, T8, and Pz) for Irreg_4–4_ in the active session, separated within participant by behavioral report into trials judged as “change” (red) and trials judged as “no change” (gray); shaded bands indicate ± s.e.m. The vertical pink line marks the nominal change point (1 s). The blue bar indicates the RM window (1000–1200 ms post-onset). **(B)** Scalp distributions of ΔRM for “change”-report and “no change”-report trials. The color scale indicates ΔRM magnitude (warm colors, positive ΔRM; cool colors, negative ΔRM). Black dots denote electrodes with ΔRM significantly greater than zero (one-sample t-test against zero, FDR-corrected *p* < 0.05). **(C)** Participant-level mean ΔRM averaged across the predefined temporal-parietal-occipital ROI for “change”-report and “no change”-report trials. *p* values above each condition indicate one-sample *t*-tests against zero; the horizontal comparison indicates the paired *t*-test between trial types.

Across example channels, “change”-report trials showed a stronger change-locked modulation around the nominal midpoint than “no change”-report trials (Fig. 5A). This difference was also apparent in scalp topographies: ΔRM was broadly positive for “change”-report trials and reached significance relative to zero at multiple electrodes (black dots), whereas “no change”-report trials showed weaker ΔRM and did not differ from zero at the ROI level (Fig. 5B–C). At the participant level, mean ΔRM averaged across the predefined temporal-parietal-occipital ROI was significantly larger for “change”-report than for “no change”-report trials (*p* = 0.03, paired *t*-test; Fig. 5C). Consistent with this, ΔRM for “change”-report trials was significantly greater than zero (*p* < 0.001, one-sample *t*-test), whereas ΔRM for “no change”-report trials was not (*p* = 0.48, one-sample *t*-test; Fig. 5C). Together, these results indicate that the predictive change-like response not only depends on stimulus statistics and task engagement (Figs. 2–3), but also tracks subjective change judgments even when the stimulus contains no physical transition.

## Discussion

Our results identify a context-defined, predictive change-like response in human EEG that can emerge in the absence of any physical stimulus transition. Behaviorally, subtle timing changes were reliably detected when click timing was fixed, but sensitivity was markedly reduced when click timing was variable (Fig. 1C). Consistent with the expanded stimulus set and the change-prevalent trial structure, participants were globally biased toward reporting “change” (Fig. 1D). At the neural level, sensory-driven change responses to physical transitions were robust in the fixed-timing condition and attenuated under variable timing (Supplementary Figs. 1–2). Crucially, during active detection the variable-timing no-change stimulus (Irreg_4–4_) elicited a midpoint-locked response that exceeded the fixed-timing no-change control (Reg_4–4_) (Fig. 2A–C), was amplified relative to passive listening (Fig. 3A–C), and remained absent in the fixed-timing no-change control across sessions (Supplementary Fig. 3). Moreover, when aligned to the nominal change point, a clear peak was observed only in the early post-change window, whereas the pre-change and late post-change windows showed no comparable response (Fig. 4). This temporal specificity is consistent with the pre-specified RM window used for ΔRM quantification (1000–1200 ms post-onset). Finally, the predictive response also tracked subjective report: within Irreg_4–4_ trials, ΔRM was larger on trials judged as “change” than on trials judged as “no change” (Fig. 5), linking the neural effect to perceptual decision state even in the absence of a physical transition. Together, these dissociations show that change-like activity can be expressed at an expected timepoint without a stimulus change, but only under a specific combination of stimulus statistics, task engagement, and a change-prevalent context.

Predictive coding frames perception as inference: the brain maintains generative models of the sensory world, compares incoming evidence against those models, and updates them when expectations fail ^2,3^. Mismatch negativity (MMN) provides a canonical population-level signature of violated expectation, reliably elicited when a discrete deviant violates an inferred regularity ^4–6^. Omission paradigms sharpen the same logic by demonstrating that time-locked cortical responses can be evoked even when an expected event is absent, indicating that prediction-related update signals can be anchored to internal timing rather than stimulus energy ^16,17^. However, most omission effects are demonstrated in rhythmic, event-based sequences, where discrete token timing and entrainment are integral to the paradigm. Our data extend this framework to continuous stimulation with a context-defined update timepoint—a fixed, task-relevant midpoint embedded within a change-prevalent design (Fig. 1A, Fig. 1D). In this regime, the response we observe is not best interpreted as a reaction to a missing sound token, but as an internally timed, prediction-error–like update signal to a missing structural transition at the expected midpoint of a continuous stream (Fig. 1A; Figs. 2–3). This shift matters because it implies that predictive machinery can be recruited not only by violations of what occurs, but also by violations of when an internally defined update is expected during ongoing input ^2,3,15^. The result also complements “phantom deviance” evidence showing that endogenous perceptual states can establish top-down predictions that yield deviance-like responses when violated ^20^. The report-dependent modulation observed in Fig. 5 is consistent with this view, suggesting that the predictive response is coupled to decision-related state under uncertainty rather than being a fixed sensory artifact.

A second key feature is gating by temporal uncertainty and task set. The predictive change-like response was selectively expressed for Irreg_4–4_ and absent for Reg_4–4_ (Fig. 2A–C; Supplementary Fig. 3), a pattern naturally captured by precision-weighted accounts in which update signals are scaled by the estimated reliability (precision) of sensory evidence relative to higher-level priors ^2,3,22^. When click timing is highly consistent (Reg_4–4_), bottom-up evidence is precise and stabilizes the internal model, suppressing spurious updating at the midpoint despite the broader task context (Fig. 2A; Supplementary Fig. 3). When timing is variable (Irreg_4–4_), sensory precision is reduced, increasing the relative influence of context-driven priors—here, an expectation that something may happen at 1 s given the experimental setting (Fig. 1A–C). Under these conditions, the absence of a transition at the expected moment can yield a temporally specific, prediction-error–like update signal expressed as a midpoint-locked change-like response (Fig. 2A–C) ^2,3,5^. The lack of any corresponding effect in Reg_4–4_, together with the window-specific response profile quantified in Fig. 4, argues against generic midpoint segmentation, slow drift, or broad state fluctuations as sufficient explanations (Supplementary Fig. 3; Fig. 4). The strong active–passive dissociation further supports a task-set contribution: active detection amplified the response (Fig. 3A–C), consistent with the idea that task engagement increases the precision of the “midpoint-change” hypothesis and boosts the gain of mismatch signals at behaviorally relevant timepoints ^3,22,23^. In passive listening, the midpoint carries less behavioral relevance, weakening the context-defined temporal prior and thereby greatly reducing the predictive response (Fig. 3A–C). Notably, the report-dependent modulation in Fig. 5 provides convergent support that the response is coupled to behavioral set rather than reflecting a purely time-locked sensory transient.

Finally, these results provide a compact empirical separation between (i) sensory-driven change responses to physical transitions (Supplementary Figs. 1–2) and (ii) a context- and task-gated predictive change-like response expressed without physical change when temporal uncertainty is high (Figs. 2–3; Supplementary Fig. 3). Notably, the effect was temporally confined to the early post-change window and was modulated by subjective change reports. More broadly, the present paradigm offers a practical assay for dissociating sensory-driven change detection from context-gated predictive updating in continuous auditory streams, and it provides a principled handle for testing how uncertainty and behavioral goals shape prediction-related neural dynamics in humans ^5,6,16,17,23^.

## Materials and methods

### Experimental procedure and participants

This study was conducted in accordance with the Declaration of Helsinki (2013) ^24^. A total of 42 participants (20 males and 22 females, mean age: 23.36 years old, standard deviation: 2.55) participated in this experiment. Participants maintained a stationary head position while listening to auditory stimuli and responding via keyboard presses. Experiments including healthy participants were approved by the Institutional Review Board (IRB-20230131-R), and informed consent was obtained from all participants.

The experiment included two sessions: a passive session followed by an active session. Eight stimulus types were presented with equal probability in each session, including regular no-change (Reg_4–4_), regular change (Reg_4–4.01_, Reg_4–4.02_, Reg_4–4.03_, and Reg_4–4.06_), irregular no-change (Irreg_4–4_), and irregular change (Irreg_4–4.06_ and Irreg_4–8_). Each stimulus was repeated a minimum of 40 times, in both sessions.

### Auditory Stimuli

This study employed transitional click trains (Fig. 1A) ^21,25,26^. Each click consisted of a 200-μs monopolar pulse. Each trial comprised a 2-s click train made of two consecutive 1-s segments, with a fixed nominal change point at 1 s. Eight stimulus types were used: regular no-change (Reg_4–4_), regular change (Reg_4–4.01_, Reg_4–4.02_, Reg_4–4.03_, and Reg_4–4.06_), irregular no-change (Irreg_4–4_), and irregular change (Irreg_4–4.06_ and Irreg_4–8_). These stimulus types were randomly interleaved with equal probability. In regular click trains, inter-click intervals (ICIs) were fixed, whereas in irregular click trains, ICIs were drawn from a Gaussian distribution with fixed mean values and satisfied the following constraints:

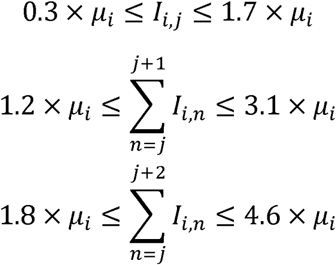

where *µ_i_* represents the mean ICI (in milliseconds) of the *i*^th^ segment in the transitional click train and *І_ij_* denotes the *j*^th^ ICI in that segment. Transitional click trains were formed by concatenating two click trains with identical or different ICIs or mean ICIs. For example, Reg_4–4.06_ consisted of a 1-s regular train with a 4-ms ICI followed seamlessly by a 1-s regular train with a 4.06-ms ICI. Stimuli were delivered through a Golden Field M23 speaker, powered by a Creative AE-7 Sound Blaster at a sampling rate of 384 kHz, and controlled via Psychtoolbox 3 in MATLAB. Sound intensity was calibrated to a constant 60 dB SPL using a ¼-inch Brüel & Kjær 4954 condenser microphone and PHOTON/RT analyzer.

### Data acquisition

Electroencephalogram (EEG) data were acquired using a 64-channel NeuroScan system (Compumedics, Australia). In practice, we only included 60 electrodes in the analyses. The ground electrode of the 64-channel NeuroScan Quick-Cap was placed between Fpz and Fz in the frontal area, while the reference electrode was placed between Cz and CPz. The EEG data were sampled at 1 kHz, and electrode placement followed the international 10-20 system protocol.

### Data Analysis

#### Preprocessing

Data analyses were performed using MATLAB R2021b (The MathWorks, Inc., Natick, MA, USA) and the FieldTrip toolbox ^27^. Monopolar referencing was applied using the default reference electrode positioned between Cz and Pz. Multichannel EEG data underwent sequential preprocessing: initial bandpass filtering (two-pass Butterworth, 0.5–40 Hz), followed by epoch extraction from −1 to 3 s relative to trial onset. Independent component analysis (ICA) removed electrooculogram (EOG) artifacts. Baseline correction subtracted the mean activity from −200 to 0 ms pre-stimulus onset per trial. Motion artifacts were evaluated via a relative threshold, flagging samples outside mean ± 3 × SD as “bad.” Trials exceeding 20% bad samples were discarded as “bad trials,” and channels with >10% bad trials were labeled “bad channels.” Bad channels were excluded first (based on all-channel data), followed by bad trials from remaining channels. Event-related potentials (ERPs) were computed by averaging epochs per experimental condition, channel, and subject. Prior to group-level analysis, ERPs were normalized by dividing each channel’s data by the standard deviation of baseline (−200 to 0 ms pre-stimulus onset) per subject to minimize inter-channel variability.

#### Permutation test

Permutation tests for time-point and channel-wise comparisons of ERP data between stimuli groups employed a two-tailed cluster-based approach via the ‘ft_timelockstatistics’ function in the FieldTrip toolbox for MATLAB. A two-tailed independent-samples t-test was conducted at each time point and channel to compute t-values, contrasting Dataset A (e.g., ERP data for Irreg_4-4_ in the passive session) and Dataset B (e.g., ERP data for Irreg_4-4_ in the active session) across 42 participants. A cluster-defining threshold of *p* < 0.05 identified significant time points, which were grouped into clusters based on temporal adjacency (consecutive significant time points) and spatial proximity (neighboring electrodes). Condition labels for Datasets A and B were randomly permuted, generating a new t-value matrix per iteration; clusters were reformed using the same *p* < 0.05 threshold. The largest cluster-level statistic (e.g., sum of t-values) from each of 1000 permutations formed the null distribution. Observed clusters were deemed significant if their statistic exceeded the 95^th^ percentile of this distribution (*p* < 0.05). Tests spanned the time window of −500 to 2500 ms relative to the transitional click train onset.

#### Quantification of EEG response

To quantify the change response, we defined the response magnitude (RM) as the root mean square (RMS) of the ERP within the predefined change-response window. For each channel, RM was calculated in the specified window (e.g., [1000, 1200] ms relative to stimulus onset, corresponding to [0, 200] ms relative to the nominal change point at 1 s). The relative response magnitude (ΔRM) was then obtained by subtracting the RMS of the baseline window [800, 1000] ms post-onset, corresponding to [−200, 0] ms pre-change. Specifically, for a channel *i*,

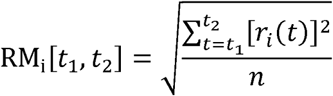

where *r_i_*(*t*) denotes the ERP of the *i^th^* channel at time *t*(ms), *n* is the number of samples within the time window [*t*_1_, *t*_2_] and RM_i_ is the response magnitude of channel *i.* The relative response magnitude was then defined as

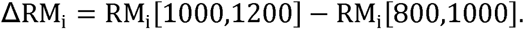

The baseline window immediately preceding the nominal change point represented the steady-state response to Train 1. To obtain an ROI-based measure of the change response, ΔRM was averaged across a predefined set of temporal-parietal-occipital channels (T7, T8, TP7, TP8, P7, P5, P3, P1, Pz, P2, P4, P6, P8, PO7, PO5, PO3, POz, PO4, PO6, PO8, O1, Oz, and O2). This ROI was chosen because the canonical change response in the transitional click-train paradigm is strongest over temporal-parietal-occipital scalp regions, with pronounced responses reported at temporal, parietal, and occipital electrodes in an independent EEG study ^21^, and the predictive change-like response observed here showed a qualitatively similar posterior distribution (Fig. 2B). Averaging within this ROI therefore provides a more sensitive summary measure while avoiding dilution of the effect by electrodes outside the main response field.

### Statistics

Behavioral comparisons of change-detection ratio between paired conditions (e.g., Irreg_4–4_ and Irreg_4–4.06_) were performed using two-tailed Wilcoxon signed-rank tests. For EEG analyses, two-tailed one-sample *t*-tests were used to test whether the average ΔRM across the predefined temporal-parietal-occipital ROI differed from zero (e.g., Fig. 2C, *p* values above each condition). Unless otherwise noted, pairwise comparisons of ΔRM between conditions were performed using two-tailed paired *t*-tests (e.g., Irreg_4–4_ in the passive versus active session).

To identify channels showing significant change responses, ΔRM was computed for each channel and subject, as shown in the topographic plots (e.g., Fig. 2B). For each channel, a two-tailed one-sample *t*-test against zero was performed across subjects, and the resulting *p* values were corrected across channels using the Benjamini-Yekutieli false discovery rate (FDR) procedure ^28^. Channels surviving correction were marked with black dots on the topographic plots.

### Data Availability

All data used in this study are available on request.

All codes applied in this study are available on a public Github repository (url: https://github.com:TOMORI233/EEGProcess.git). All source data supporting the findings of this study are publicly available at Zenodo under the following DOI: https://doi.org/10.5281/zenodo.15795285 ^29^.

## Author Contributions

Conceptualization: X.Y.; Methodology: H.X., W.W., Y.Y., X.Y., B.Z.; Software: H.X., W.W.; Formal analysis: H.X., W.W.; Investigation: H.X., W.W., Y.Y., I.M.; Data curation: H.X., W.W., Y.Y.; Visualization: H.X., W.W.; Writing—original draft: H.X., W.W., Y.Y., X.Y.; Writing—review & editing: H.X., W.W., Y.Y., I.M., Z.D., M.M., B.Z., X.Y.; Resources: X.Y.; Supervision: X.Y.; Funding acquisition: X.Y.

## Acknowledgments

This work was supported by Brain Science and Brain-like Intelligence Technology––National Science and Technology Major Project (2022ZD0204600 and 2022ZD0204800) (to X.Y.); National Natural Science Foundation of China 32571216 and 32171044 (to X.Y.);

