## Supplementary Figures 1-3 for "Expectation triggers a change-like EEG response without acoustic change"

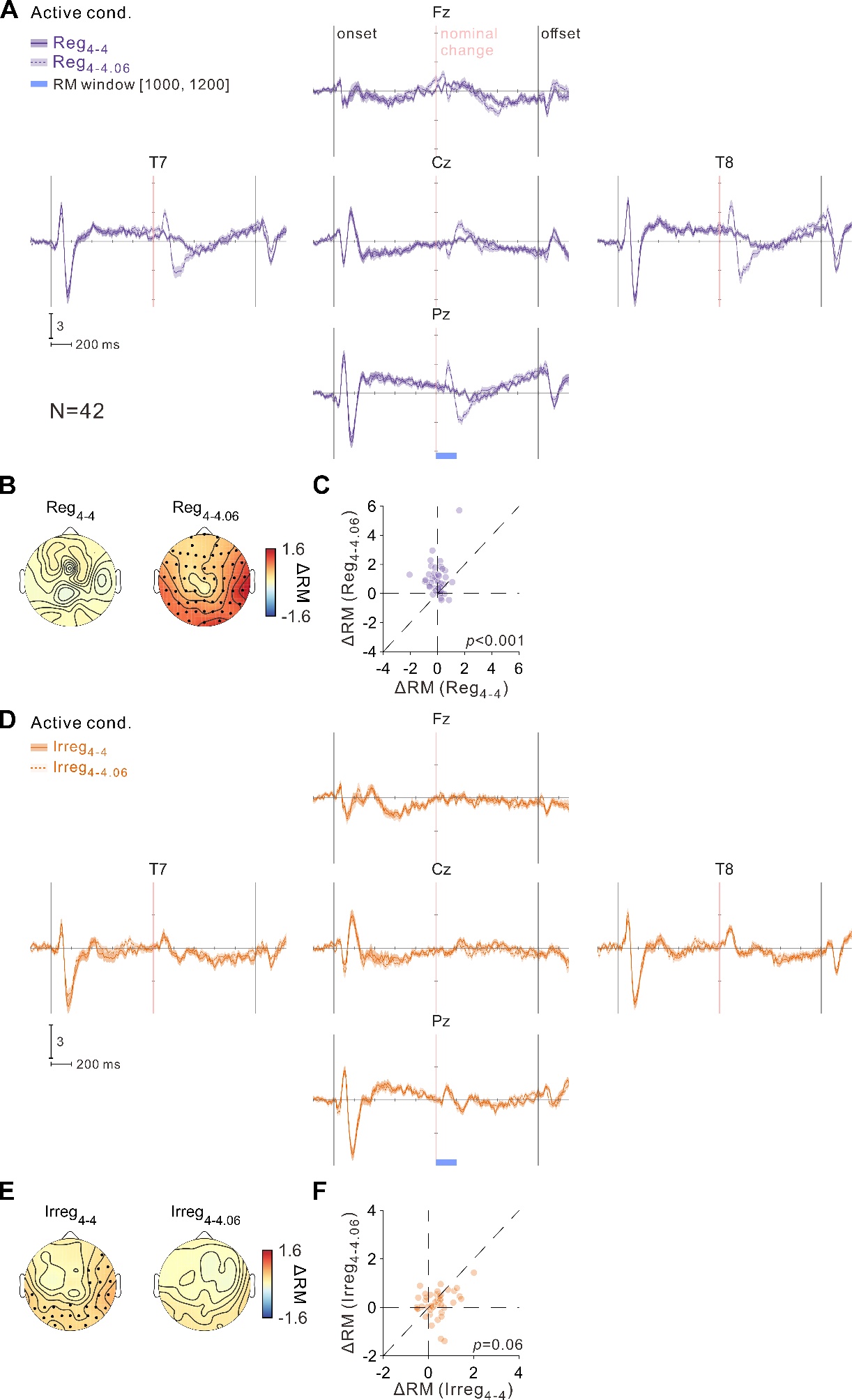

**Supplementary Figure 1. Sensory-driven change responses to physical transitions in the active session.** **(A–C)** Comparison of **Reg_4–4_** and **Reg_4–4.06_** in the **active** session. **(A)** Grand-average ERPs (N = 42) from five example channels (Fz, T7, Cz, T8, and Pz); shaded bands indicate ± s.e.m. **Solid purple** indicates Reg_4–4_ and **dashed purple** indicates Reg_4–4.06_. The vertical pink line marks the nominal change point (1 s). The blue bar indicates the RM window (1000–1200 ms post-onset). **(B)** Scalp distributions of ΔRM for Reg_4–4_ and Reg_4–4.06_. The color scale indicates ΔRM magnitude (warm colors, positive ΔRM; cool colors, negative ΔRM). Black dots denote electrodes with ΔRM significantly greater than zero (one-sample *t*-test against zero, FDR-corrected *p* < 0.05). **(C)** Participant-level comparison of mean ΔRM averaged across the predefined temporal-parietal-occipital ROI between Reg_4–4_ and Reg_4–4.06_; each point represents one participant, the diagonal line indicates equality, and the *p* value denotes the paired t-test between conditions. **(D–F)** Same analyses as in **(A–C)**, but comparing Irreg_4–4_ and Irreg_4–4.06_ in the active session. **(D)** Grand-average ERPs; **solid orange** indicates Irreg_4–4_ and **dashed orange** indicates Irreg_4–4.06_ (± s.e.m.). **(E)** Scalp distributions of ΔRM (same color scale and significance criteria as in B). **(F)** Participant-level comparison of mean ΔRM averaged across the predefined temporal-parietal-occipital ROI between Irreg_4–4_ and Irreg_4–4.06_ (paired *t*-test).

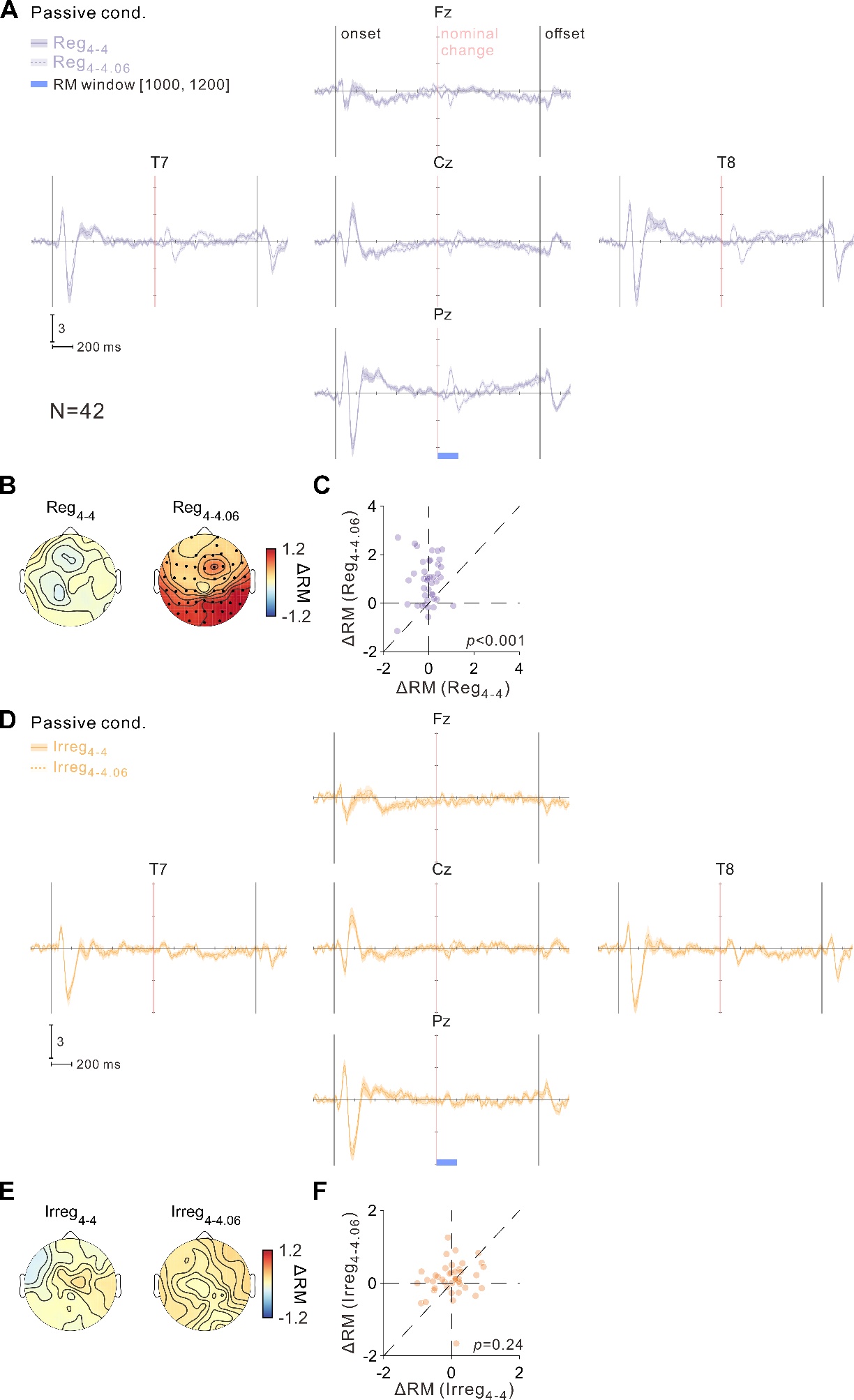

**Supplementary Figure 2. Sensory-driven change responses to physical transitions in the passive session.** Same analyses as in Supplementary Fig. 1, but for the passive session. **(A–C)** Comparison of Reg_4–4_ and Reg_4–4.06_. **(A)** Grand-average ERPs (N = 42) from five example channels (Fz, T7, Cz, T8, and Pz); shaded bands indicate ± s.e.m. Solid light purple indicates Reg_4–4_ and dashed light purple indicates Reg_4–4.06_. The vertical pink line marks the nominal change point (1 s). The blue bar indicates the RM window (1000–1200 ms post-onset). **(B)** Scalp distributions of ΔRM for Reg_4–4_ and Reg_4–4.06_. The color scale indicates ΔRM magnitude (warm colors, positive ΔRM; cool colors, negative ΔRM). Black dots denote electrodes with ΔRM significantly greater than zero (one-sample *t*-test against zero, FDR-corrected *p* < 0.05). **(C)** Participant-level comparison of mean ΔRM averaged across the predefined temporal-parietal-occipital ROI; each point represents one participant, the diagonal line indicates equality, and the *p* value denotes the paired t-test between conditions. **(D–F)** Same analyses as in **(A–C)**, but comparing Irreg_4–4_ and Irreg_4–4.06_ in the passive session. **(D)** Grand-average ERPs; solid light orange indicates Irreg_4–4_ and dashed light orange indicates Irreg_4–4.06_ (± s.e.m.). **(E)** Scalp distributions of ΔRM (same color scale and significance criteria as in **B**). **(F)** Participant-level comparison of mean ΔRM averaged across the predefined temporal-parietal-occipital ROI between Irreg_4–4_ and Irreg_4–4.06_ (paired *t*-test).

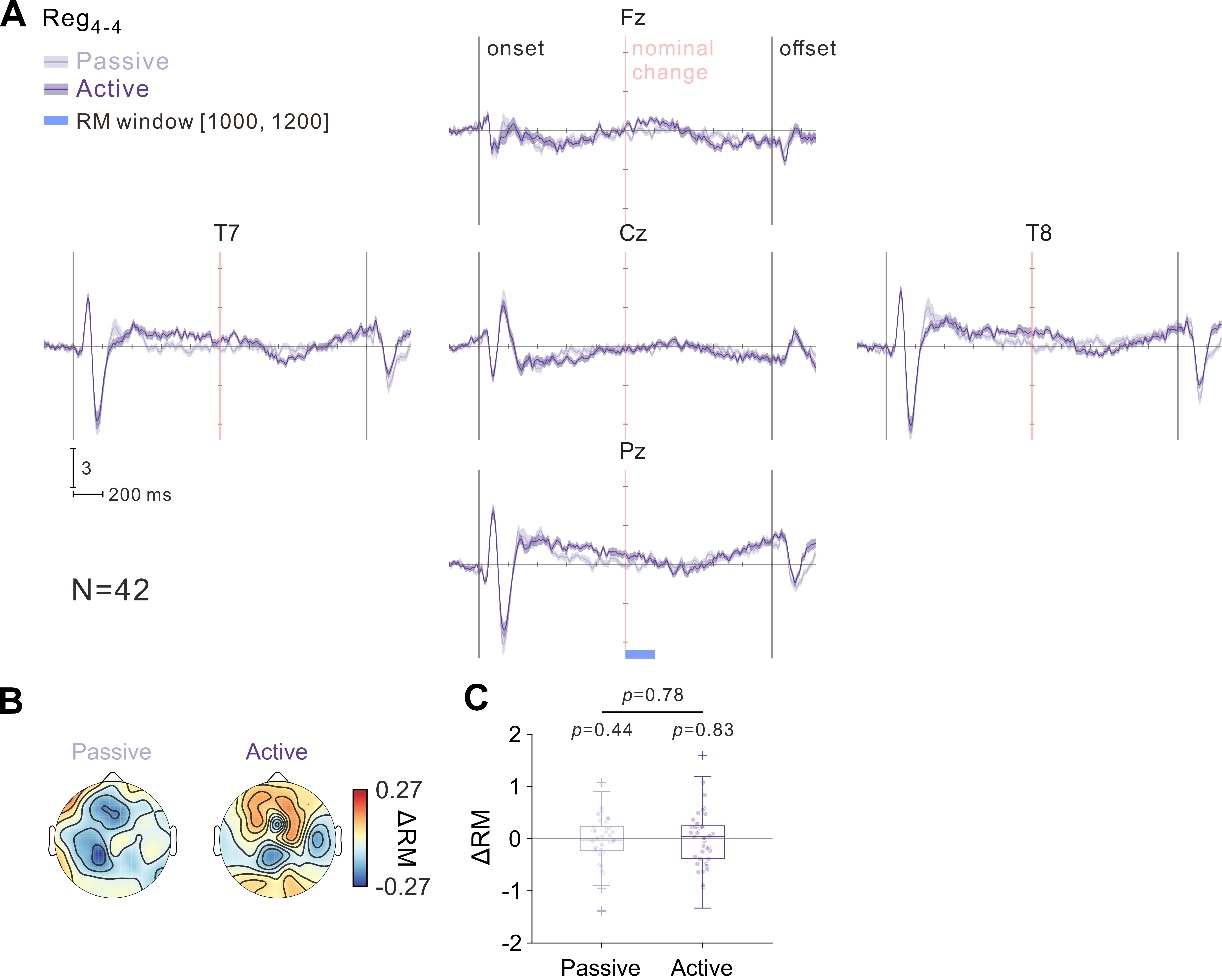

**Supplementary Figure 3. Absence of predictive change-like responses in the fixed-timing no-change control.** **(A)** Grand-average ERPs (N = 42) from five example channels (Fz, T7, Cz, T8, and Pz) for Reg_4–4_ in the passive and active sessions; shaded bands indicate ± s.e.m. Light purple indicates passive and dark purple indicates active. The vertical pink line marks the nominal change point (1 s), and the blue bar indicates the RM window (1000–1200 ms post-onset). **(B)** Scalp distributions of ΔRM for Reg_4–4_ in the passive and active sessions. The color scale indicates ΔRM magnitude (warm colors, positive ΔRM; cool colors, negative ΔRM). No electrode showed ΔRM significantly greater than zero (one-sample *t*-test against zero, FDR-corrected *p* < 0.05). **(C)** Participant-level mean ΔRM averaged across the predefined temporal-parietal-occipital ROI in the passive and active sessions. *p* values above each condition indicate one-sample *t*-tests against zero; the horizontal comparison indicates the paired *t*-test between sessions.
